# Multiple variants of the blast fungus effector AVR-Pik bind the HMA domain of the rice protein OsHIPP19 with high affinity

**DOI:** 10.1101/2020.12.01.403451

**Authors:** Josephine H. R. Maidment, Marina Franceschetti, Abbas Maqbool, Hiromasa Saitoh, Chatchawan Jantasuriyarat, Sophien Kamoun, Ryohei Terauchi, Mark J. Banfield

## Abstract

Microbial plant pathogens secrete effector proteins which manipulate the host to promote infection. Effectors can be recognised by plant intracellular nucleotide-binding leucine-rich repeat (NLR) receptors, initiating an immune response. The AVR-Pik effector from the rice blast fungus *Magnaporthe oryzae* is recognised by a pair of rice NLR receptors, Pik-1 and Pik-2. Pik-1 contains a non-canonical integrated heavy metal-associated (HMA) domain, which directly binds AVR-Pik to activate plant defences. Non-canonical integrated domains are widespread in plant NLRs and are thought to resemble the host target of the recognised effector. AVR-Pik interacts with specific rice HMA domain-containing proteins, namely heavy metal-associated isoprenylated plant proteins (HIPPs) and heavy metal-associated plant proteins (HPPs). Here, we define the biochemical and structural basis of the interaction between AVR-Pik and OsHIPP19, and compare the interaction with the HMA domain of Pik-1. Using analytical gel filtration and surface plasmon resonance, we show that multiple AVR-Pik variants, including the stealthy variants AVR-PikC and AVR-PikF which do not interact with any characterised Pik-1 alleles, bind to OsHIPP19 with nanomolar affinity. The crystal structure of OsHIPP19 in complex with AVR-PikF reveals differences at the interface that underpin high-affinity binding of OsHIPP19-HMA to a wider set of AVR-Pik variants than achieved by the integrated HMA domain of Pik-1. Our results provide a foundation for engineering the HMA domain of Pik-1 to extend binding to currently unrecognised AVR-Pik variants and expand disease resistance in rice to divergent pathogen strains.

## Introduction

Phytopathogens constrain crop production and threaten global food security. The filamentous ascomycete fungus *Magnaporthe oryzae* is the causative agent of rice blast disease, which annually destroys enough rice to feed upwards of 200 million people for a year (1,2). *M. oryzae* is found in all major rice-growing regions around the world (3), and severe epidemics cause total crop loss (2,4). In addition to rice, strains of *M. oryzae* can infect and cause blast disease on other staple food crops such as wheat, barley and millet, and various wild grass species (5,6).

During infection, phytopathogens deliver effector proteins to the plant apoplast and to the inside of host cells. These effectors function in diverse ways to suppress immunity and manipulate endogenous processes to create favourable conditions for colonisation. While many *M. oryzae* effector candidates have been identified and cloned (7), only a few have been functionally characterised (8–13). For most *M. oryzae* effectors described to date, the mechanism by which they promote fungal virulence remains unknown.

The presence of some effectors inside host cells can be detected by specific nucleotide-binding, leucine-rich repeat domain containing proteins (NLRs) (14). NLRs are intracellular immune receptors which instigate defensive signalling pathways following perception of a specific effector protein, either through direct interaction with the effector, or through monitoring the state of an intermediate protein (15–17). The canonical NLR structure comprises a C-terminal leucine-rich repeat domain, a central nucleotide-binding NB-ARC domain, and an N-terminal coiled-coil (CC or CC_R_ (18)) or Toll/interleukin-1 receptor (TIR) domain (19–21). Recent studies have identified non-canonical domains in multiple NLR proteins from different plant species (22–24). These integrated domains (IDs) are thought to have their evolutionary origins in the host targets of the effector (25,26). By acting as an effector substrate or interactor, they enable the NLR to detect the presence of the effector and trigger host immunity.

The paired rice CC-NLRs Pik-1 and Pik-2 are genetically linked and cooperate to trigger immunity in response to the *M. oryzae* effector AVR-Pik (27–29). Pik-1 contains an integrated heavy metal-associated (HMA) domain between the canonical CC and NB-ARC domains (figure 3a) (29). Previous work demonstrated that detection of AVR-Pik is through direct binding of the effector to the integrated HMA domain of Pik-1 (29). Five Pik-1 alleles have been identified and cloned (27,30–32), with most polymorphisms located in and around the integrated HMA domain. The integrated HMA domain of Pik-1 is thought to resemble the host target of AVR-Pik. While it has been speculated that AVR-Pik may target rice proteins containing an HMA domain (33), until recently (34) these targets have remained undefined.

Six AVR-Pik variants (A-F) have been identified to date, differing in just five amino acid positions (28,35–37). Each of these polymorphic amino acids are located at the interface with Pik-1-HMA (29), indicating that they are adaptive. While *M. oryzae* strains carrying AVR-PikD trigger immune responses in rice lines containing Pikp, Pikm, Pikh or Pik*, strains carrying either AVR-PikE or AVR-PikA only elicit a response in rice lines with Pikm or Pikh (28,29,36,38). No naturally occurring Pik alleles respond to the stealthy effectors AVR-PikC and AVR-PikF.

In an independent study (34), rice proteins targeted by the AVR-PikD effector were identified by a yeast 2-hybrid screen. Four small heavy metal-associated domain-containing (sHMA) proteins were identified as interactors of AVR-PikD. OsHIPP19 (LOC_Os04g39350) and OsHIPP20 (LOC_Os04g39010) are members of the <u>h</u>eavy metal-associated isoprenylated <u>p</u>lant <u>p</u>rotein (HIPP) family (39,40), while OsHPP03 (LOC_Os02g37290) and OsHPP04 (LOC_Os02g37300) belong to the <u>h</u>eavy metal-associated <u>p</u>lant <u>p</u>rotein (HPP) family. All four proteins contain a N-terminal HMA domain, and OsHIPP19 and OsHIPP20 have a CααX isoprenylation motif (-CSIM) at their C-termini (figure 1).

**Figure 1.**
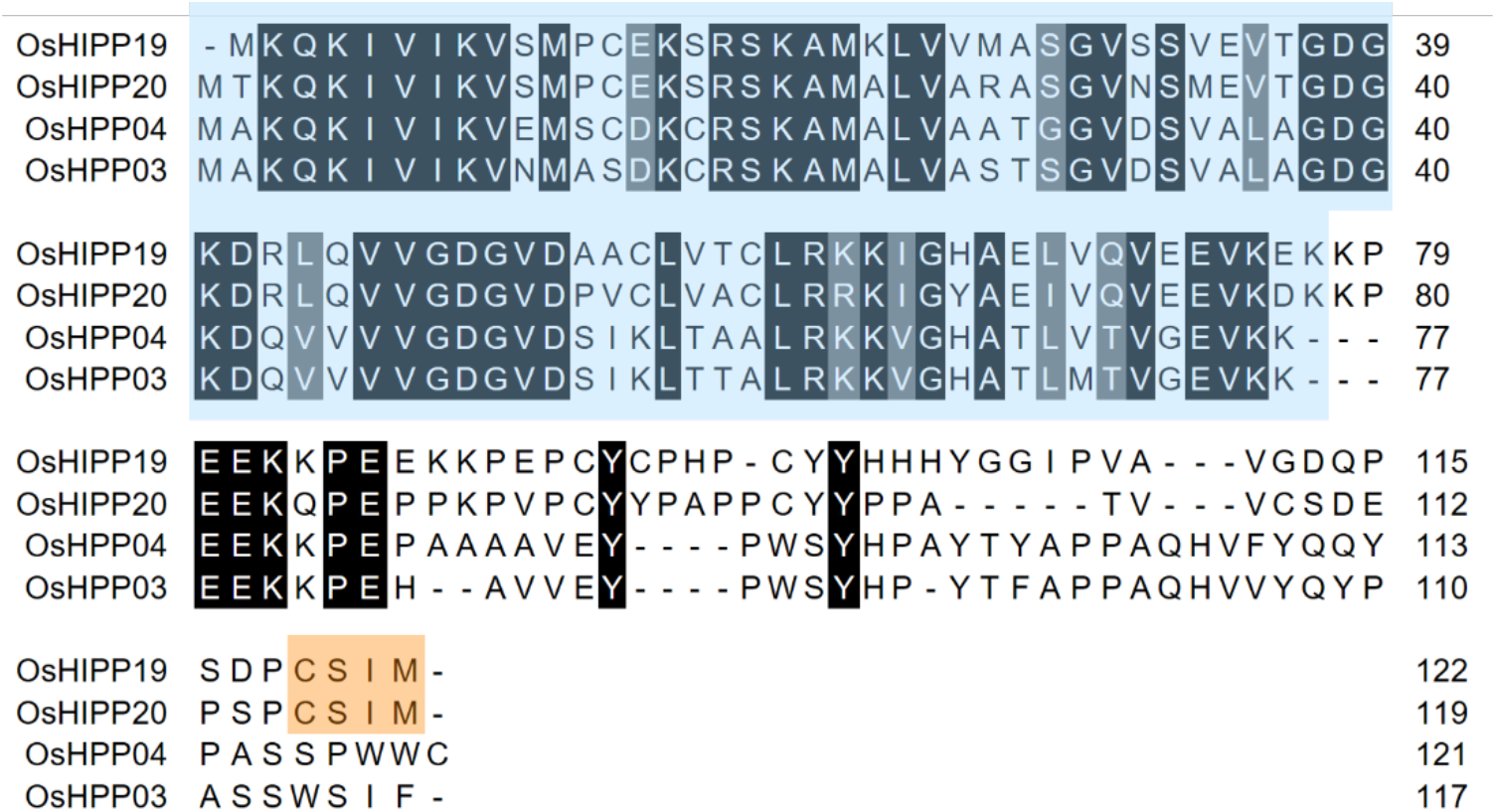
Amino acid sequence alignment of the AVR-PikD interactors identified by yeast two-hybrid (34), OsHIPP19 (LOC_Os04g39350), OsHIPP20 (LOC_Os04g39010), OsHPP03 (LOC_Os02g37290) and OsHPP04 (LOC_Os02g37300). The alignment was produced with Clustal Omega and coloured using BoxShade. The HMA domain is highlighted in blue, and the isoprenylation motif in OsHIPP19 and OsHIPP20 is highlighted in orange.

Here, taking a biochemical and structural approach, we characterise the interaction between AVR-Pik and OsHIPP19. We demonstrate that AVR-PikD interacts with the HMA domain of OsHIPP19 with high affinity, and that the interaction between AVR-PikD and the HMA domain of OsHIPP19 is tighter than the interactions between AVR-PikD and the integrated HMA domains of Pik-1 alleles. Further, we show that the tight interaction with OsHIPP19 is not unique to AVR-PikD, but is shared with other AVR-Pik variants. Finally, we present the crystal structure of AVR-PikF in complex with the HMA domain of OsHIPP19, revealing differences between the effector-binding surfaces of OsHIPP19 and that of the NLR protein Pikm-1. Our findings reveal the biochemical and structural basis of a *M. oryzae* effector protein’s interaction with a putative virulence-associated host target, and present opportunities for target-guided engineering of NLR integrated domains to extend their recognition profile.

## Results

### AVR-PikD binds to the HMA domain of OsHIPP19 with nanomolar affinity

In an independent study, four small HMA domain-containing proteins were found to interact with AVR-PikD by yeast 2-hybrid analysis (34). Of these, we selected OsHIPP19 for further study as it was identified most frequently in the initial yeast 2-hybrid screen (34). Based on the well-characterised interaction of AVR-PikD with the HMA domain of Pik-1 (29,41,42), we hypothesised that AVR-PikD would interact with the HMA domain of OsHIPP19. We defined the HMA domain of OsHIPP19 (OsHIPP19-HMA hereafter) as amino acids 1-77 inclusive. This domain was expressed and purified from *E. coli* (see Materials and Methods) and used in all subsequent work in vitro. Intact mass spectrometry confirmed a molecular mass of 8323 Da for the purified protein, which exactly matched the theoretical mass.

First, we used analytical gel filtration to qualitatively test for interaction between AVR-PikD and OsHIPP19-HMA. When combined, the two proteins co-eluted from the column, at an earlier elution volume than observed for either protein alone, demonstrating that AVR-PikD and OsHIPP19-HMA form a complex in vitro (figure 2a). The *Phytophthora infestans* effector PexRD54, which targets the autophagy-related protein ATG8 in its host (*Solanum tuberosum*) (43,44) was used as a negative control, and did not co-elute with OsHIPP19-HMA, nor was a shift in elution volume observed for PexRD54 in the presence of OsHIPP19 (supplementary figure 1).

**Figure 2.**
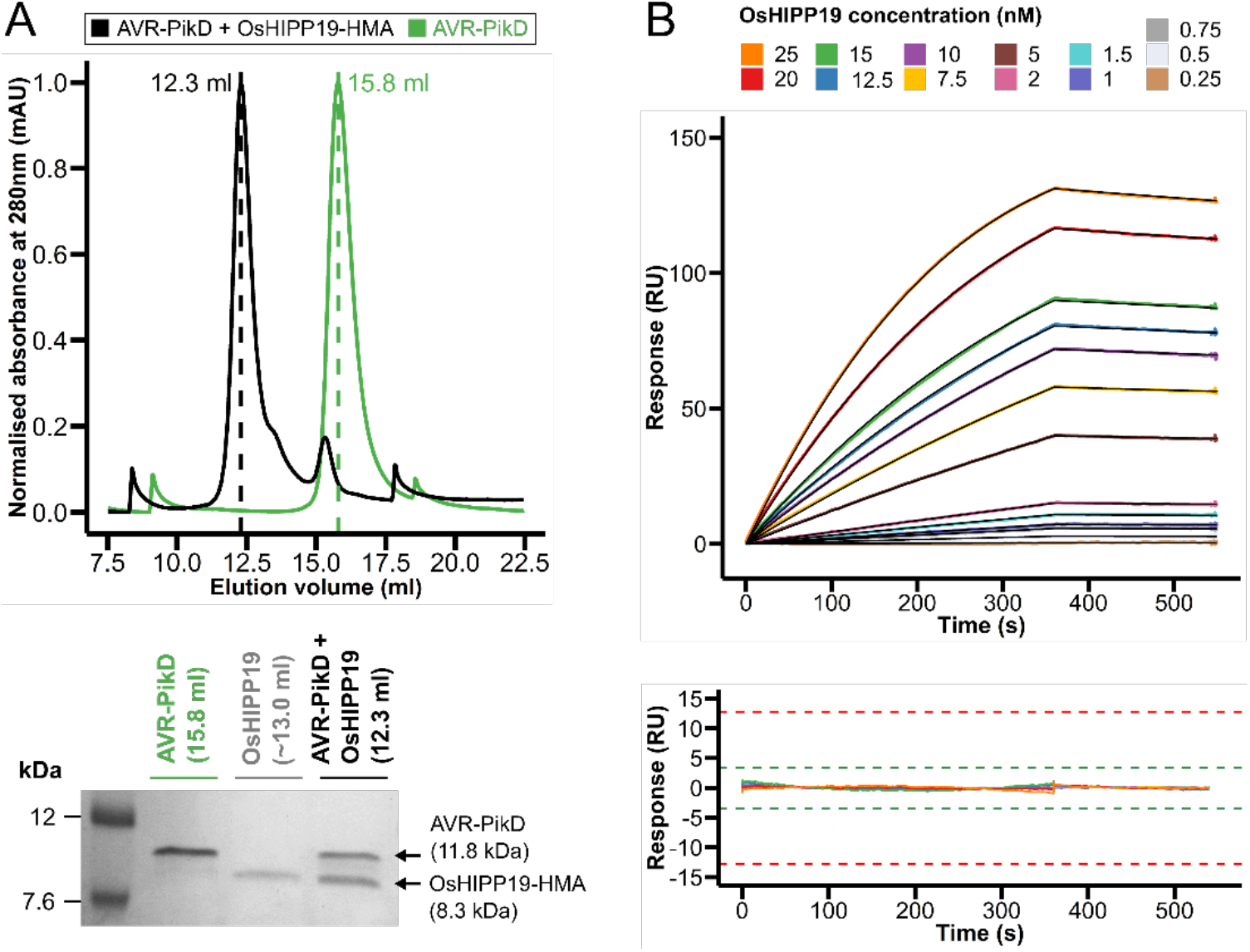
AVR-PikD interacts with the HMA domain of OsHIPP19 with nanomolar affinity *in vitro*. A) Normalised analytical gel filtration traces for AVR-PikD alone (green) and AVR-PikD with OsHIPP19-HMA (black). The SDS-PAGE gel shows fractions from the peak elution volumes of each sample, demonstrating that AVR-PikD and OsHIPP19 co-elute. OsHIPP19-HMA absorbs UV light at 280nm very poorly (molar extinction coefficient of 360 cm^-1^M^-1^), so no peak is visible for OsHIPP19-HMA. B) Multicycle kinetics data from surface plasmon resonance (coloured lines) and 1-to-1 binding model fitted to the data (black lines) with the residuals plotted below. Green and red acceptance thresholds are determined by the Biacore T100 evaluation software.

To investigate the affinity of the interaction between AVR-PikD and OsHIPP19-HMA, we used surface plasmon resonance (SPR). AVR-PikD was immobilised on a Ni^2+^-NTA chip via a non-cleavable 6xHis tag at the C-terminus of the protein. OsHIPP19-HMA, at a range of concentrations between 0.25 nM and 25 nM, was flowed over the chip. A 1:1 binding model was used to fit the data (figure 2b) and to estimate the rate constants k_a_ and k_d_ for the interaction (supplementary table 1), from which the equilibrium dissociation constant, *K_D_*, could be calculated (*K_D_* = k_d_/k_a_). The *K_D_* value obtained from the data shown in figure 2b was 0.7 nM. Two further replicates were performed with similar results (*K_D_* = 0.6nM and *K_D_* = 0.9 nM). We conclude that AVR-PikD binds to OsHIPP19 with nanomolar affinity.

### AVR-PikD binds to the HMA domain of OsHIPP19 with higher affinity than to the integrated HMA domains of Pikp-1 and Pikm-1

The HMA domain of OsHIPP19 shares 51% amino acid sequence identity with the integrated HMA domains of both Pikp-1 and Pikm-1 (figure 3a, figure 3b). Previous work has shown that AVR-PikD binds to the integrated HMA domains of Pikp-1 and Pikm-1 with nanomolar affinity (29,42). To compare the binding of AVR-PikD to the integrated HMA domains with its binding to OsHIPP19-HMA, we again used surface plasmon resonance (SPR). AVR-PikD was immobilised on a Ni^2+^-NTA chip via a C-terminal 6xHis tag. Three different concentrations of each HMA domain (2 nM, 5 nM and 20 nM) were flowed over the chip, and the binding (R_obs_, measured in response units (RU)) was recorded. The R_obs_ for each HMA was then expressed as a percentage of the maximum theoretical response (R_max_) that would be obtained if each immobilised molecule of AVR-PikD was bound to the HMA domain, referred to as %R_max_. Mutating glutamate-230 of Pikp-HMA to arginine was predicted to disrupt the interaction of the HMA domain with AVR-PikD, and therefore Pikp^E230R^-HMA was used as a negative control. Consistent with previous data, AVR-PikD interacted with the HMA domains of Pikp-1 and Pikm-1 with similar apparent affinity (figure 3c, supplementary figure 2). Interestingly, AVR-PikD bound to OsHIPP19-HMA with higher apparent affinity (larger %R_max_) than to the HMA domains of either Pikp-1 and Pikm-1 (figure 3c, supplementary figure 2) at each of the three concentrations tested.

**Figure 3.**
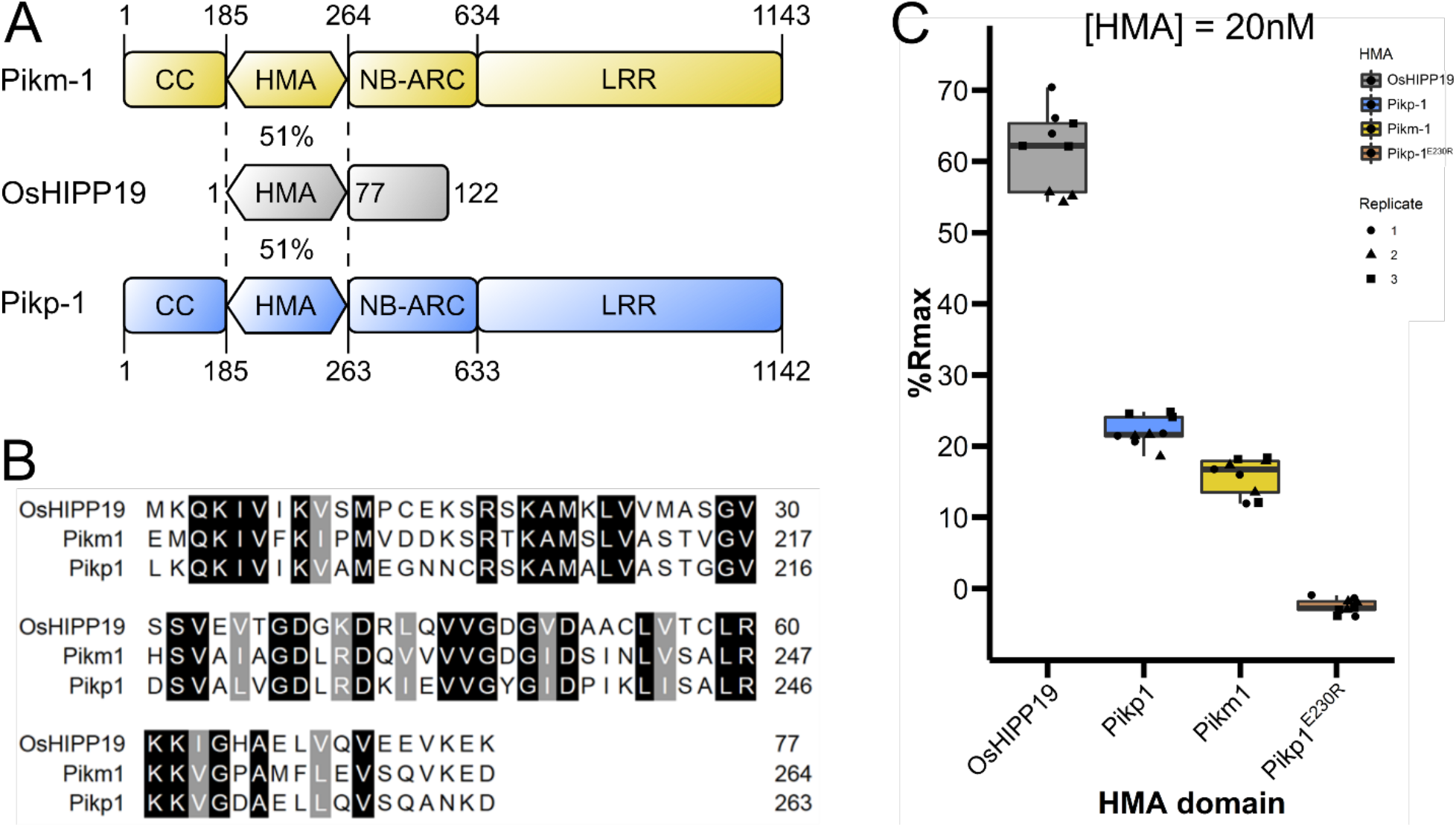
AVR-PikD interacts with the HMA domain of OsHIPP19 with higher affinity than with the integrated HMA domains of Pikp-1 or Pikm-1. A) Schematic representation of Pikm-1, Pikp-1 and OsHIPP19. OsHIPP19-HMA shares 51% sequence identity with both Pikp-HMA and Pikm-HMA. B) Amino acid sequence alignment of the HMA domains of OsHIPP19, Pikp-1 and Pikm-1. The alignment was produced with Clustal Omega and coloured using BoxShade. C) Boxplots showing the %R_max_ observed for the interactions between AVR-PikD and each of the HMA domains. %R_max_ is the percentage of the theoretical maximum response, assuming a 1:1 HMA:effector binding model for OsHIPP19-HMA and Pikm-HMA, and a 2:1 binding model for Pikp-HMA and Pikp^E230R^-HMA. The centre line of the box represents the median and the box limits are the upper and lower quartiles. The whiskers extend to the smallest value within Q_1_ – 1.5x the interquartile range (IQR) and the largest value within Q_3_ + 1.5x IQR. Individual data points are represented as black shapes. The experiment was repeated three times, with each experiment consisting of three technical replicates. Plots were produced using the ggplot2 package (Wickham, 2016) in R (R Core Development Team, 2018).

### Additional AVR-Pik variants interact with the HMA domain of OsHIPP19

The five AVR-Pik variants A, C, D, E and F differ in only 5 amino acid positions (supplementary figure 3). Previous work has shown that only a subset of AVR-Pik variants interact with the HMA domains of Pikp-1 and Pikm-1 (29,42). Notably, neither AVR-PikC nor AVR-PikF interact with the HMA domains of Pikp-1 and Pikm-1, and therefore neither variant triggers Pik-mediated immunity in plants. We hypothesised that AVR-PikC and AVR-PikF may have evolved to avoid detection by the plant immune system, but still interact with their host target (OsHIPP19) to execute their virulence function.

To test this hypothesis, we used analytical gel filtration to qualitatively assess whether each of the AVR-Pik variants forms a complex with OsHIPP19 in vitro. We found that each of the AVR-Pik variants, including AVR-PikC and AVR-PikF, formed a complex with OsHIPP19-HMA (figure 4, supplementary figure 4). To investigate whether the amino acid polymorphisms between the different AVR-Pik variants influence their affinity for OsHIPP19, we determined the equilibrium dissociation constants for the interactions between OsHIPP19-HMA and AVR-PikC/AVR-PikF by SPR, as described earlier for AVR-PikD. The *K_D_* values obtained for these interactions over three replicates were within a similar nanomolar range (1.1-1.9 nM for AVR-PikC, and 0.8-1.0 nM for AVR-PikF) (figure 5a and 5b, supplementary table 1), indicating that AVR-PikC and AVR-PikF also bind OsHIPP19 with similar nanomolar affinity.

**Figure 4.**
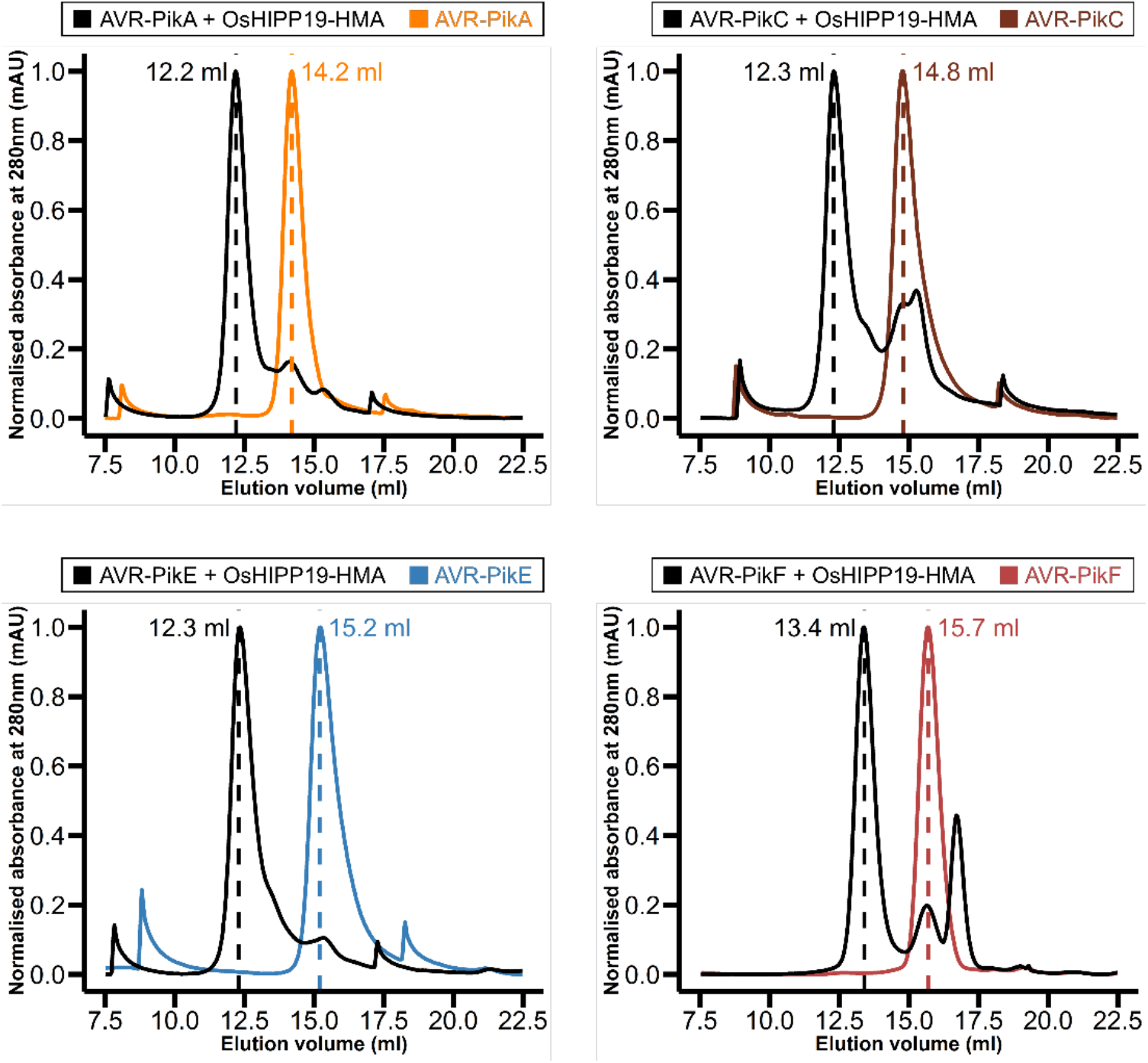
All AVR-Pik alleles interact with the HMA domain of OsHIPP19. Normalised analytical gel filtration traces for each effector allele alone (coloured) and with OsHIPP19-HMA (black). The peak shift observed when each effector is combined with OsHIPP19-HMA indicates complex formation. OsHIPP19-HMA absorbs UV light at 280 nm very poorly (molar extinction coefficient of 360 cm^-1^M^-1^), so no peak is visible for OsHIPP19-HMA.

**Figure 5.**
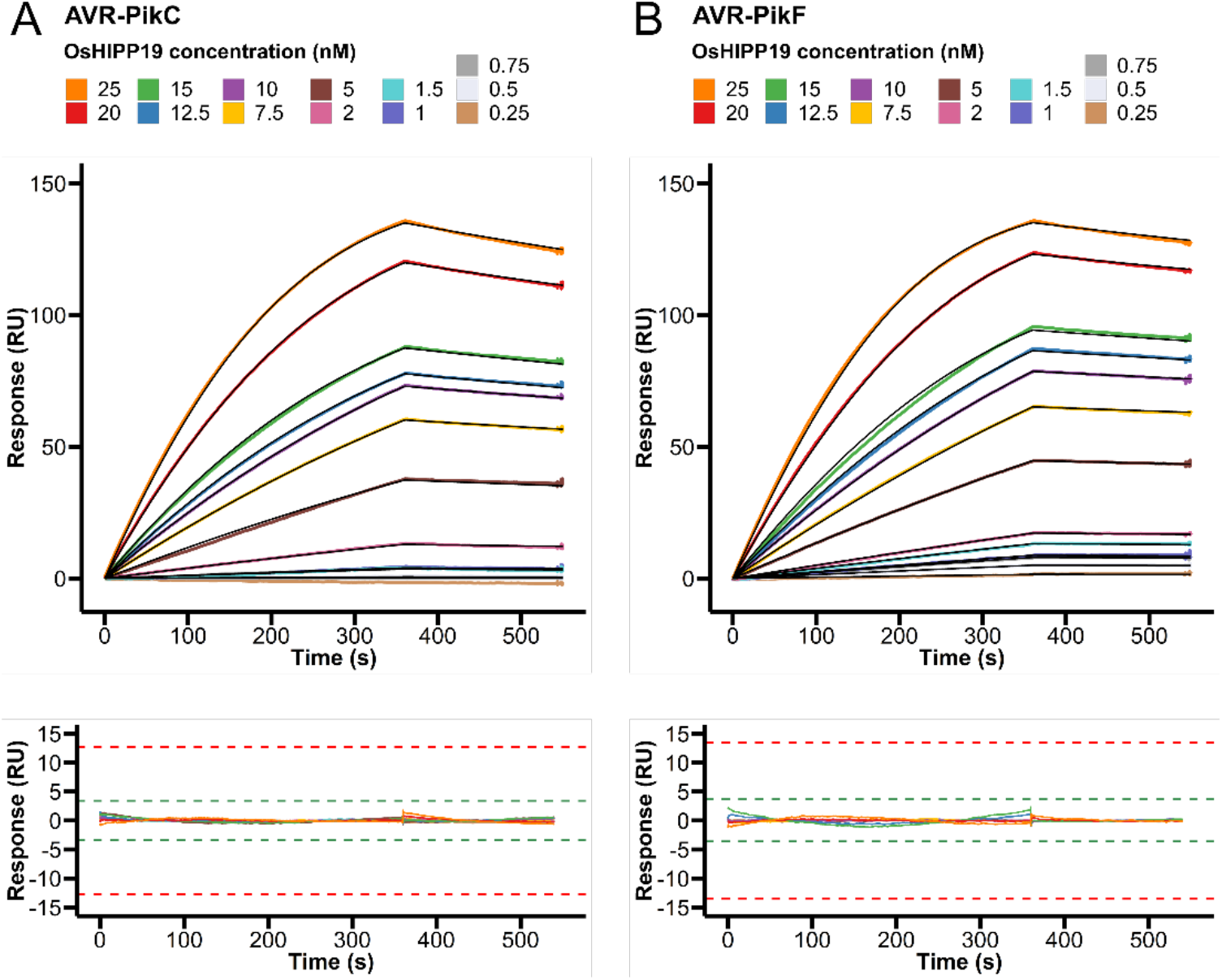
AVR-PikC and AVR-PikF interact with the HMA domain of OsHIPP19-HMA with nanomolar affinity. Multicycle kinetics data from surface plasmon resonance experiments for the interaction between OsHIPP19-HMA and A) AVR-PikC or B) AVR-PikF. Experimental data is shown as coloured lines, and 1-to-1 binding model fitted to the data overlaid (black lines). The residuals are plotted below; green and red acceptance thresholds are determined by the Biacore T100 evaluation software.

### Crystal structure of AVR-PikF in complex with OsHIPP19

Next, we defined the structural basis of interaction between OsHIPP19 and AVR-Pik effectors. As AVR-PikC and AVR-PikF do not bind any of the integrated HMA domains identified to date, their interaction with the HMA domain of OsHIPP19 was of particular interest. Both AVR-PikC and AVR-PikF were each co-expressed with OsHIPP19-HMA in *E. coli* and purified (see Materials and Methods for full details). Crystals of the complex between OsHIPP19-HMA and AVR-PikF were obtained in several conditions in the commercial Morpheus screen (Molecular Dimensions). The OsHIPP19-HMA/AVR-PikC complex did not crystallise in any of the commercial screens trialled. We therefore focused on OsHIPP19-HMA/AVR-PikF. Having confirmed the presence of both proteins in the purified complex by intact mass spectrometry (peaks in the spectrum at 8323 Da and 10839 Da exactly matched the theoretical masses for OsHIPP19 and AVR-PikF (after subtracting 2 Da for the presence of an expected disulphide bond), respectively), we then obtained X-ray diffraction data to 1.9Å resolution. The structure of the OsHIPP19-HMA/AVR-PikF complex was solved by molecular replacement (see Materials and Methods), using the crystal structure of a monomer of Pikp-HMA/AVR-PikD (PDB accession no. 5A6W) as a model. X-ray data collection, refinement, and validation statistics are shown in table 1.

**Table 1.** Data collection and refinement statistics for the AVR-PikF/OsHIPP19-HMA complex.

| Data collection statistics |  |
| --- | --- |
| Wavelength (Å) | 0.976250 |
| Space group | P 2<sub>1</sub> 2<sub>1</sub> 2<sub>1</sub> |
| Cell dimensions *a*, *b*, *c* (Å) | 29.78, 53.78, 98.03 |
| Resolution (Å)* | 98.03-1.90 (1.94-1.90) |
| R<sub>merge</sub> (%) | 12.9 (112.9) |
| *I* / $\sigma$ *I* | 12.4 (1.9) |
| Completeness (%) | 100 (100) |
| Unique reflections | 13077 (811) |
| Redundancy | 12.6 (11.1) |
| CC(1/2) (%) | 99.9 (91.0) |
| Refinement and model statistics |  |
| Resolution (Å)* | 98.03-1.90 (1.95-1.90) |
| R<sub>work</sub>/R<sub>free</sub> (%) | 19.6/23.3 (27.5/27.1) |
| No. atoms |  |
| Protein | 2416 |
| Water | 67 |
| B-factors |  |
| Protein | 38.1 |
| Water | 37.4 |
| R.m.s deviations |  |
| Bond lengths (Å) | 0.015 |
| Bond angles (°) | 1.661 |
| Ramachandran plot (%)** |  |
| Favoured | 98.0 |
| Allowed | 2.0 |
| Outliers | 0 |
| MolProbity Score | 1.39 (98<sup>th</sup> percentile) |

\* The highest resolution shell is shown in parentheses.
\*\* As calculated by MolProbity

As anticipated, the HMA domain of OsHIPP19 adopts the well-characterised HMA fold (Pfam: PF00403) consisting of a four-stranded antiparallel β-sheet and two α-helices arranged in an α-β sandwich. The loop between β1 and α1 containing the degenerate metal-binding motif (MPCEKS) is poorly defined in the electron density, and OsHIPP19^Glu14-Lys15^ could not be positioned.

Overall, the structure of AVR-PikF is very similar to previously determined structures of other AVR-Pik variants, and comprises a core six-stranded β-sandwich, conserved among the MAX effectors (45), with an N-terminal extension (AVR-PikF^Arg31-Pro52^). A disulphide bond between AVR-PikF^Cys54^ and AVR-PikF^Cys70^ stabilises the β-sandwich structure.

OsHIPP19-HMA and AVR-PikF form a 1:1 complex. The position of AVR-PikF relative to the HMA domain of OsHIPP19 is similar to the previously determined structures of AVR-Pik effectors in complex with integrated Pik-HMA domains (figure 6) (29,42). AVR-PikF only differs from AVR-PikA by a single polymorphism at position 78, and the structures of the OsHIPP19-HMA/AVR-PikF complex and Pikm-HMA/AVR-PikA complex are similar. The RMSD, as calculated in Coot (46) using secondary structure matching, between the HMA domains is 0.826Å using 74 residues. The RMSD between the AVR-Pik effectors is 0.437Å using 81 residues. The overall RMSD between the two complexes is 0.71Å using 155 residues.

**Figure 6.**
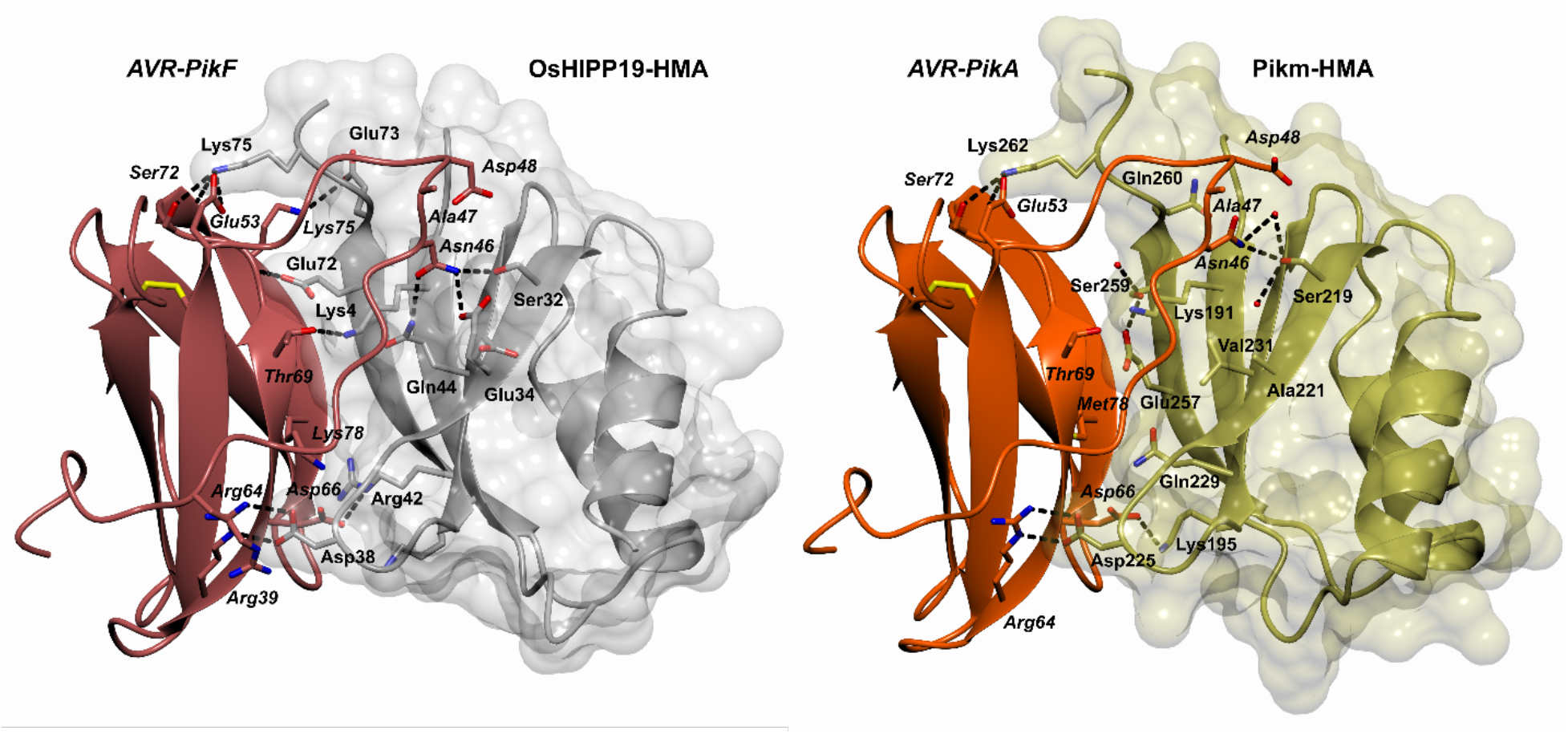
Comparison of the crystal structure of AVR-PikF in complex with the HMA domain of OsHIPP19 (7B1I) with the previously published crystal structure of AVR-PikA in complex with the HMA domain of Pikm-1 (6FUD, (42)). The structures of AVR-PikF and OsHIPP19-HMA are represented as red and grey ribbons, respectively, with the molecular surface of OsHIPP19-HMA also shown. The structures of AVR-PikA and Pikm-HMA are represented as orange and gold ribbons, respectively, with the molecular surface of Pikm-HMA also shown. The side chains of amino acids of interest are displayed as cylinders. Hydrogen bonds are represented by dashed lines.

The interface between OsHIPP19-HMA and AVR-PikF is extensive, burying 23.1% and 19.5% of the total accessible surface area of the HMA domain (1068.5Å^2^) and effector (1022.0Å^2^) respectively. The total interface area (sum of the buried surface area of each component divided by two) for the OsHIPP19-HMA/AVR-PikF complex is 1045.3Å^2^, larger than the total interface area of the Pikm-HMA/AVR-PikA complex (918.3Å^2^) (42), and indeed any of the Pik-HMA/AVR-Pik complexes determined to date. Interface analysis parameters determined by QtPISA are shown in supplementary table 2.

### Differences over three interfaces, but particularly at interface 3, underpin the higher affinity of OsHIPP19 for AVR-Pik relative to the integrated Pikm-1 HMA domain

Previous analysis of the interface between AVR-PikA and Pikm-HMA revealed three main regions, numbered 1 to 3, which contribute to the interaction between the two proteins. Similarly, three distinct regions can be identified in the complex of OsHIPP19-HMA/AVR-PikF. Differences between OsHIPP19-HMA and Pikm-HMA at each of these three interfaces may contribute to the differences in specificity and affinity of the interactions between the HMA domains and AVR-Pik variants.

Interface 1 of AVR-PikA/Pikm was characterised by a weak (3.5Å) hydrogen bond between the sidechain of Pikm^Lys191^ and the main-chain carbonyl group of AVR-PikA^Thr69^, and a hydrophobic interface contributed by Pikm^Met189^ (42). In the AVR-PikF/OsHIPP19 complex, the hydrophobic interface is absent, however the lysine residue is conserved and forms a second hydrogen bond (2.9Å) with the side-chain of AVR-PikF^Thr69^ (figure 6).

In both complexes, interface 2 involves residues from β2 and β3 of the HMA domain (Pikm^Ser219-Val233^ and OsHIPP19^Ser31-Val46^), which interact with residues in β2 and the N-terminal extension of AVR-PikF (including the polymorphic residues 46, 47 and 48). AVR-PikA and AVR-PikF share the asparagine-alanine-aspartate (NAD) triad in the polymorphic positions 46, 47 and 48. The interactions between these residues and residues in β2 and β3 of the HMA domain underpin the differential recognition of AVR-PikD, AVR-PikE, and AVR-PikA by Pikm (42). The side-chain of AVR-PikA^Asn46^ forms a single hydrogen bond with Pikm^Ser219^. By contrast, the side-chain of AVR-PikF^Asn46^ is rotated and forms two hydrogen bonds with the side-chains of OsHIPP19^Ser32^ and OsHIPP19^Gln44^ (figure 7a). Also located to interface 2, AVR-PikF^Asp66^ forms hydrogen bonds with the main-chain amide group of OsHIPP19^Asp38^ and side-chain of OsHIPP19^Arg42^, differing from the AVR-PikA/Pikm structure where a single hydrogen bond is formed between AVR-PikA^Asp66^ and Pikm^Lys195^. The side-chain of Pikm^Asp225^ forms two hydrogen bonds with the side-chain of AVR-PikA^Arg64^; the aspartate is conserved in OsHIPP19 (OsHIPP19^Asp38^) and forms similar interactions with AVR-PikF^Arg64^ (figure 6).

**Figure 7.**
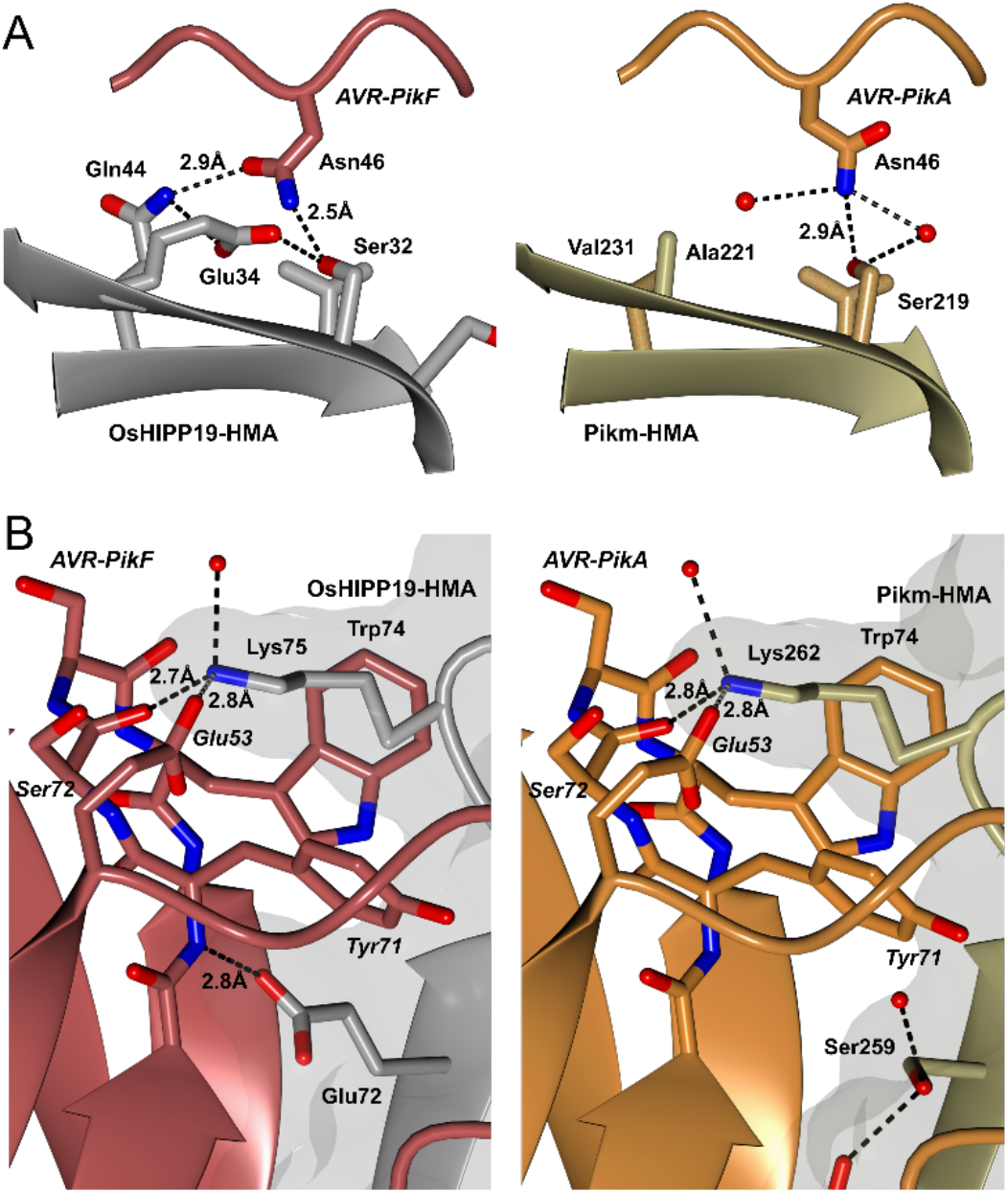
Comparison of the binding interfaces in the crystal structure of AVR-PikF in complex with the HMA domain of OsHIPP19 (7B1I) with the previously published crystal structure of AVR-PikA in complex with the HMA domain of Pikm-1 (6FUD, (42)). The structures of AVR-PikF and OsHIPP19-HMA are represented as red and grey ribbons, respectively, with the molecular surface of OsHIPP19-HMA also shown. The structures of AVR-PikA and Pikm-HMA are represented as orange and gold ribbons, respectively, with the molecular surface of Pikm-HMA also shown. The side chains of amino acids of interest are displayed as cylinders. Hydrogen bonds are represented by dashed lines. A) AVR-PikF^Asn46^ is rotated and forms an additional hydrogen bond with OsHIPP19-HMA compared to AVR-PikA^Asn46^ bound to Pikm-HMA. The side-chain of OsHIPP19^Glu34^ exists in two alternate conformations, both supported by the electron density. For clarity, only the relevant conformation is shown here. B) OsHIPP19^Glu72^ forms an additional hydrogen bond with the main chain of AVR-PikF.

Interface 3 comprises residues from β4 of the HMA domain extending to the C-terminus of the protein (Pikm^Met254-Asp264^ and OsHIPP19^Glu67-Glu76^). In the complex between Pikm-HMA/AVR-PikA, this interface is defined by main-chain hydrogen bonding between β4 of the HMA domain and β3 of AVR-PikA and, notably, the positioning of Pikm^Lys272^ into a pocket on the surface of the effector. The conserved OsHIPP19^Lys75^ binds into a similar pocket on the surface of AVR-PikF, formed by AVR-PikF^Glu53^, AVR-PikF^Tyr71^, AVR-PikF^Ser72^ and AVR-PikF^Trp74^. In addition to main-chain hydrogen bonding between β4 of OsHIPP19-HMA and β3 of AVR-PikF, additional hydrogen bonds are contributed by OsHIPP19^Glu72^ and OsHIPP19^Glu73^. Strikingly, the side-chain of OsHIPP19^Glu72^ (a serine in the corresponding position in Pikm-HMA) forms a hydrogen bond (2.8Å) with the main-chain amide group of AVR-PikF^Tyr71^ (figure 7b). In addition, the side-chain of OsHIPP19^Glu73^ (a glutamine in the corresponding position in Pikm-HMA) forms a salt bridge interaction with the side chain of AVR-PikF^Lys75^. Overall, hydrogen bonding between the HMA domain and the effector at interface 3 is more extensive in the OsHIPP19-HMA/AVR-PikF complex than in the Pikm-HMA/AVR-PikA complex, which likely contributes to the difference in binding affinity.

### Polymorphic residue AVR-PikF^Lys78^ is located at the binding interface with OsHIPP19

The single difference between AVR-PikA, which binds to Pikm-1 to trigger plant immunity, and AVR-PikF, which does not, is the residue at position 78 (methionine in AVR-PikA and lysine in AVR-PikF). The polymorphic AVR-PikF^Lys78^ is positioned at the binding interface with OsHIPP19 and is well defined in the electron density. Interestingly, the sidechain of the lysine adopts an unusual conformation (determined by MolProbity with reference to lysine sidechains in deposited structures in the protein data bank) (supplementary figure 5). We suggest that while this residue is sufficient to disrupt the interaction between the integrated HMA domains and AVR-PikF, increased intermolecular interactions between OsHIPP19 and AVR-PikF, as described earlier, compensate for the disruptive influence of AVR-PikF^Lys78^ and maintain the interaction between the two proteins.

## Discussion

Recognition of the *M. oryzae* effector AVR-Pik is mediated by the paired rice NLR proteins Pik-1/Pik-2. AVR-Pik directly interacts with an integrated heavy metal-associated (HMA) domain in Pik-1. However, the stealthy variants AVR-PikC and AVR-PikF avoid binding to the integrated Pik-1 HMA domain and evade plant defences. Here we show that all AVR-Pik variants, including AVR-PikC and AVR-PikF, interact with the HMA domain of rice heavy-metal associated isoprenylated plant protein 19 (OsHIPP19), a putative virulence target, with nanomolar affinity (figure 8). We observe that AVR-Pik variants interact with OsHIPP19-HMA with higher apparent affinity than with the integrated Pik-HMA domains. By solving the crystal structure of AVR-PikF bound to OsHIPP19-HMA and comparing it to the previously published structures of AVR-Pik effectors bound to Pikp-HMA or Pikm-HMA, we identify differences in the binding interfaces that underpin the increased affinity and broader specificity of the OsHIPP19-HMA/AVR-Pik interaction.

**Figure 8.**
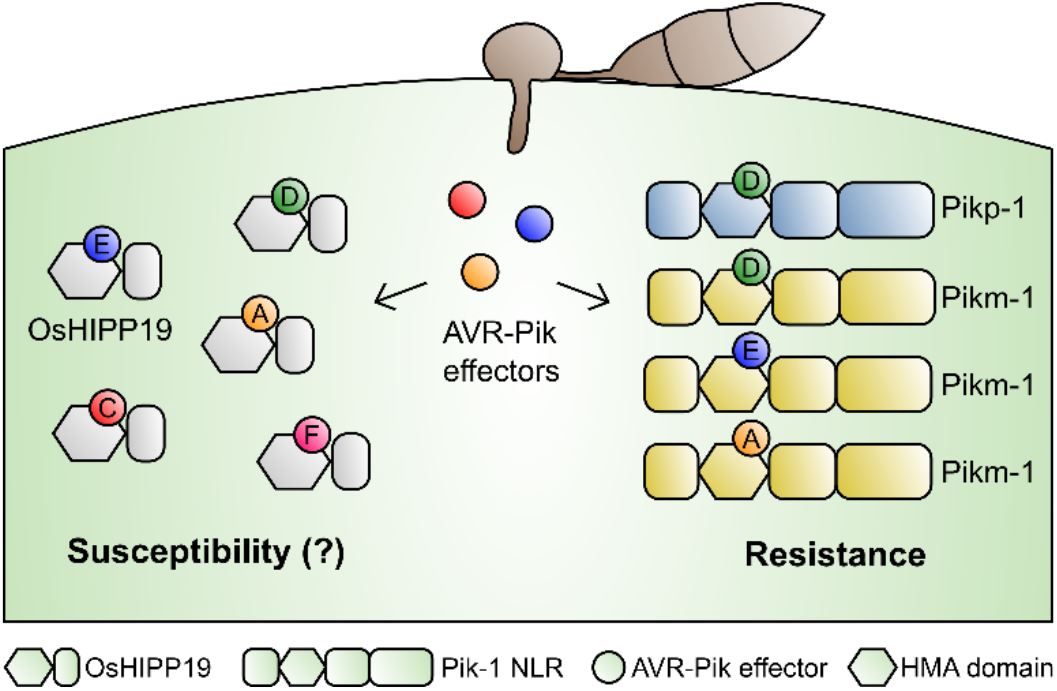
Schematic representation of the interactions between AVR-Pik effector alleles and the HMA domains of OsHIPP19 and Pik-1 NLR proteins.

### AVR-Pik binds OsHIPP19-HMA with higher affinity than integrated Pik-1 HMA domains

The identification of diverse protein domains integrated into the core structure of NLRs has provided new insights into the molecular mechanisms by which NLR proteins detect the presence of effectors. However, little is known about how effectors interact with host virulence-associated targets that are related to these integrated domains. Direct binding of AVR-Pik to the integrated HMA domain of Pik-1 is necessary for activation of immunity by the co-operative NLR proteins Pik-1 and Pik-2. Similarly, the rice NLR pair Pia (also known as RGA5/RGA4) detects the presence of AVR-Pia or AVR1-CO39 via a direct interaction between the effector and the HMA domain at the C-terminus of the Pia sensor NLR (RGA5) (22,33,47). The integrated domain hypothesis proposes that these domains have their origins in the host virulence targets of the effector (25). As effector binding triggers immune signalling, we might expect the integrated HMA domains of Pikp-1 and Pikm-1 to bind the effector more tightly than the HMA domain of the putative virulence target OsHIPP19, as their only expected function is to bait the effector. Somewhat surprisingly, our results show that AVR-PikD interacts with OsHIPP19-HMA with higher affinity than Pikp-HMA or Pikm-HMA.

It is important to note that the HMA domains are being studied in isolation, without the biological context of the full-length protein. Other Pik-1 domains may provide additional contacts with the effector which increase the affinity of the receptor for AVR-PikD. Furthermore, the molecular details of the mechanism by which Pik-1 and Pik-2 cooperate to initiate immune signalling remain unknown; Pik-2 may also influence the affinity of the interaction between AVR-PikD and Pik-1.

### HIPPs and HPPs may be plant immunity hubs, and are targeted by multiple pathogen proteins

Although AVR-PikD interacts with Pikp-HMA and Pikm-HMA with lower apparent affinity than with OsHIPP19-HMA, the affinity of the interaction is sufficient to trigger immune signalling and disease resistance. Therefore, higher affinity binding of the integrated HMA domain to the effector may not provide any additional evolutionary benefit. Furthermore, it is has been suggested that AVR-Pia and AVR1-CO39 may also interact with rice HMA domain-containing proteins (33). Studies of the interactomes of effectors from different phytopathogens have identified common host targets, proposed to be immunity-related “hubs” which are targeted by diverse pathogens to achieve infection (48). The movement protein of potato mop top virus interacts with NbHIPP26 to promote long-distance viral movement (49), and AtHIPP27 is reported to be a susceptibility factor for infection by the beet cyst nematode *Heterodera schachtii* (50). There may be more as-yet unidentified effectors, produced by diverse pathogens, which target HIPPs or HPPs to promote infection. It is also tempting to speculate that the Pik-1/Pik-2 NLR pair may recognise effectors outside the AVR-Pik allelic series, trading high affinity binding to AVR-Pik for binding to other effectors. The NLR pair RRS1/RPS4 recognises the structurally diverse effectors AvrRps4 from *Pseudomonas syringae pv. pisi* and PopP2 from *Ralstonia solanacearum* via an integrated WRKY domain at the C-terminus of RRS1 (51–53), highlighting how the incorporation of a domain targeted by multiple effectors into an NLR can lead to immunity to diverse pathogens.

### AVR-Pik binds to OsHIPP19-HMA via a similar interface to the integrated HMA domain of Pik-1

As the integrated domain hypothesis proposes that these domains have their origins in the host virulence targets of the effector, it would be expected that the effector binds in a similar manner to both its virulence target and to the integrated domain of the NLR protein. Consistent with this, we observe that the overall binding interface between AVR-PikF and the HMA domain of OsHIPP19 is very similar to that between AVR-Pik and the integrated HMA domains of Pikp-1 and Pikm-1 (RMSD for the complex is 0.71Å using 155 amino acids). We hypothesise that other effector/NLR-ID interfaces would likewise mimic the interface between the effector and its host target.

### The stealthy effector variants AVR-PikC and AVR-PikF bind to OsHIPP19-HMA, but not to Pik-1 HMA domains

The Ala67Asp mutation that distinguishes AVR-PikC from AVR-PikE, and the Met78Lys mutation that distinguishes AVR-PikF from AVR-PikA, are both located at the interface of the effector with the HMA domain (37,38). These mutations appear to be adaptive (37,38,42) and sufficient to prevent interaction with the integrated HMA domains of the various Pik-1 alleles. Here we show that these mutations do not preclude interaction between the effector and the HMA domain of OsHIPP19. The crystal structure of the complex between OsHIPP19/AVR-PikF reveals that the sidechain of AVR-PikF^Lys78^ adopts an unusual conformation, likely imposed by a steric clash of more favourable conformations with OsHIPP19-HMA. This suggests that the OsHIPP19/AVR-PikF interaction is maintained by additional intermolecular contacts between the proteins, particularly at interface 3, rather than compensatory mutations in the HMA domain to accommodate the AVR-PikF^Lys78^ sidechain. AVR-PikC contains an aspartate in position 67, but is otherwise identical to AVR-PikE, which has alanine in this position. We have previously shown that the side chain of AVR-PikC^Asp67^ disrupts hydrogen bonding between the effector and the HMA domain at interface 2 (38). Pikp-HMA does not interact with AVR-PikC, however Pikh-HMA, which differs in a single amino acid at interface 3, is able to interact with this effector variant (38). This further demonstrates that unfavourable interactions can be overcome by compensatory mutations at other interfaces. This has broad implications for engineering protein-protein interactions, highlighting that compensatory mutations need not be physically located next to unfavourable amino acids.

Aside from integrated domains, other NLR regions, particularly LRR domains, can directly interact with pathogen effectors (54). In these cases, it is unlikely that the interface between the NLR and effectors directly mimics the interaction with virulence-associated targets. This has implications for the evolution of recognition. It suggests effectors can escape recognition via mutation at one interface without impacting the effector’s virulence-associated activity. Recognition via integrated domains may be more robust, as mutation in an effector that prevents recognition may also reduce the protein’s disease-promoting activity. However, as is exemplified by the AVR-PikC and AVR-PikF effector variants, it can still be overcome without compromising the interaction between the effector and its target. Engineering integrated domains to more closely resemble the host target of an effector may provide more durable resistance, particularly where the virulence function of the effector is necessary for disease.

### Future opportunities for engineering integrated domains to extend the recognition capabilities of NLRs

The emergence of an allelic series of AVR-Pik effectors may have been driven by the deployment of rice varieties carrying Pik-1 alleles in agriculture (55–57). AVR-PikC and AVR-PikF avoid binding to the integrated HMA domain of Pik-1, and *M. oryzae* isolates carrying AVR-PikC and AVR-PikF are virulent on Pik-containing rice lines. The spread of isolates containing these novel variants represents a threat to global rice production (37). Understanding the interaction between AVR-PikF and the HMA domain of OsHIPP19 will guide future efforts to engineer a Pik-1 receptor with an HMA domain resembling that of OsHIPP19, to which these stealthy variants bind. Such a variant may be capable of delivering disease resistance to *M. oryzae* strains carrying AVR-PikC and AVR-PikF.

## Materials & Methods

### Protein production and purification

AVR-Pik effector proteins were produced with a cleavable Nt SUMO tag and a non-cleavable Ct His tag from pOPIN-E (58) in *E. coli* SHuffle (59) and purified as previously described (29). HMA domains were produced with a cleavable Nt His-MBP tag from pOPIN-M in *E. coli* SHuffle. Pikp-HMA and Pikm-HMA were produced and purified as previously described (29). For production of OsHIPP19-HMA, inoculated 1L cell cultures were grown in autoinduction media (60) at 30°C. 1 mM TCEP was included in the buffer for the final gel filtration of OsHIPP19-HMA. Protein concentration was determined using a Direct Detect^®^ Infrared Spectrometer (Millipore Sigma).

### Analytical gel filtration

Experiments were conducted at 4°C using a Superdex 75 10/300 GL column (GE Healthcare) equilibrated in running buffer (20 mM HEPES pH 7.5, 150 mM NaCl, 1 mM TCEP). To investigate whether the effector and OsHIPP19-HMA form a complex, the two proteins were combined in a 1:1 molar ratio and incubated on ice for 1 hour prior to analysis. Each protein was also analysed alone, at a concentration equivalent to that present in the complex. For each experiment, 100 μl protein was injected at a flow rate of 0.5 ml/min, and 500 μl fractions were collected for analysis by SDS-PAGE.

### Surface plasmon resonance

Surface plasmon resonance was carried out using a Biacore T100 (GE Healthcare) at 25°C and at a flow rate of 30 μl/minute. The running buffer was 20 mM HEPES pH 7.5, 860 mM NaCl and 0.1%(v/v) Tween^®^20. Flow cell (FC) 2 of an NTA chip (GE Healthcare) was activated with 30 μl 0.5 mM NiCl2. 30 μl of the 6xHis-tagged effector (the ligand) was immobilised on FC2 to give a response between 200 and 400 RU. The HMA domain (the analyte) was then flowed over both FC1 and FC2 for 360 s, followed by a dissociation time of 180 s. The NTA chip was regenerated after each cycle with 30 μl 0.35 M EDTA pH 8.0.

The background response from FC1 (non-specific binding of the HMA domain to the chip) was subtracted from the response from FC2. For %R_max_ experiments, the binding response (R_obs_) was measured immediately prior to the end of injection and expressed as a percentage of the theoretical maximum response (R_max_) assuming a 1:1 HMA:effector binding model for OsHIPP19-HMA and Pikm-HMA, and a 2:1 HMA:effector binding model for Pikp-HMA and Pikp^E230R^-HMA, calculated as follows:

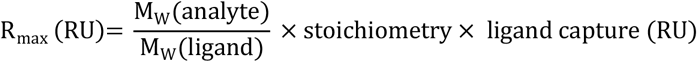

For kinetics analysis, a 1:1 binding model was applied to the data. The R_max_ was fitted locally to reflect the regeneration of the chip and subsequent recapture of Ni^2+^ and effector between cycles. The resulting estimates for the k_a_ and k_d_ values were used to calculate the reported *K_D_*.

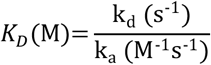

### Co-purification and crystallisation of OsHIPP19-HMA/AVR-PikF

MBP-OsHIPP19-HMA (pOPIN-A) and 6xHis-SUMO-AVR-PikF (pOPIN-S3C) were co-expressed in E. coli SHuffle cells. 8 x 1 L of cell culture was grown in auto-induction media, containing carbenicillin and kanamycin (final concentration of 100 μg/ml and 30 μg/ml, respectively) to select for the plasmids, at 30°C for 20 hours. Cells were pelleted by centrifugation (5663 x *g* for 7 minutes at 4°C) and resuspended in lysis buffer (50 mM Tris-HCl pH 8.0, 50 mM glycine, 0.5 M NaCl, 20 mM imidazole, 5% (v/v) glycerol, 1 cOmplete™ EDTA-free protease inhibitor cocktail tablet (Roche) per 50 ml buffer). Resuspended cells were lysed by sonication with a VibraCell sonicator (SONICS), and whole cell lysate was clarified by centrifugation at 38,724 x *g* for 30 minutes at 4°C. An AKTA Xpress (GE Healthcare) system was used to carry out a two-step purification at 4°C. The clarified cell lysate was first injected onto a 5 ml Ni^2+^-NTA column (GE Healthcare). The MBP-OsHIPP19-HMA/6xHis-SUMO-AVR-PikF complex was step-eluted with elution buffer (50 mM Tris-HCl pH 8.0, 50 mM glycine, 0.5 M NaCl, 500 mM imidazole, 5% (v/v) glycerol), and applied to a Superdex 75 26/600 gel filtration column equilibrated in running buffer (20 mM HEPES pH 7.5 and 150 mM NaCl). MBP and 6xHis-SUMO tags were cleaved by incubation with 3C protease (1 μg protease per mg of fusion protein) at 4°C overnight, and removed by passing the sample through a 5 ml Ni^2+^-NTA column connected to a 5 ml dextrin sepharose (MBPTrap) column (GE Healthcare), both equilibrated in lysis buffer. Fractions containing the OsHIPP19-HMA/AVR-PikF complex were concentrated and injected onto a Superdex 75 26/600 column equilibrated in running buffer + 1 mM TCEP. Protein concentration was determined using a Direct Detect^®^ Infrared Spectrometer (Millipore Sigma).

Crystallisation trials were set up in 96 well plates using an Oryx Nano robot (Douglas Instruments). 0.3 μl of 11 mg/ml protein was combined with 0.3 μl reservoir solution from the commercial Morpheus screen (Molecular Dimensions). Crystals appeared in several conditions. For X-ray diffraction, a crystal from condition D9 (0.12 M Alcohols [0.2 M 1,6-Hexanediol; 0.2 M 1-Butanol; 0.2 M 1,2-Propanediol; 0.2 M 2-Propanol; 0.2 M 1,4-Butanediol; 0.2 M 1,3-Propanediol], 0.1 M Buffer System 3 pH 8.5 [1 M Tris (base); BICINE], 50% v/v Precipitant Mix 1 [40% v/v PEG 500* MME; 20% w/v PEG 20000]) was cryoprotected in mother liquor, mounted in a loop, and frozen in liquid nitrogen.

### Data collection and model refinement

X-ray diffraction data were collected on beamline i03 at the Diamond Light Source, UK. 3600 images were collected with an exposure time of 0.01 s and oscillation of 0.1°. The diffraction images were indexed and scaled using the autoPROC pipeline (61). The scaled but unmerged data file was passed to AIMLESS (62) for data reduction. The structure was solved by molecular replacement, carried out with PHASER (63) using a monomer of Pikp-HMA in complex with AVR-PikD (PDB accession: 5A6W) as a model. OsHIPP19-HMA and AVR-PikF were built into the data using BUCCANEER (64). Iterative cycles of manual adjustment, refinement and validation were carried out using COOT (46) and REFMAC (65) through CCP4i2 (66). Final validation was performed by MolProbity (67). qtPISA (68) was used for interface analysis. Structure figures were produced using the CCP4mg molecular graphics software (69).

## Data availability

The protein structure of the complex between OsHIPP19-HMA and AVR-PikF, and the data used to derive this, have been deposited at the PDB with accession number 7B1I.

## Acknowledgements

We thank the Diamond Light Source (beamline I03) for access to X-ray data collection facilities. We thank Professor David Lawson and Dr Clare Stevenson (JIC X-ray Crystallography/Biophysical Analysis Platform) for help with X-ray data collection and SPR, and Andrew Davies and Phil Robinson from JIC Scientific Photography. We also thank Dr Freya Varden for constructive discussions. This work was supported by the UKRI Biotechnology and Biological Sciences Research Council (BBSRC) Norwich Research Park Biosciences Doctoral Training Partnership, UK [grant BB/M011216/1]; the UKRI BBSRC, UK [grants BB/P012574, BB/M02198X]; the European Research Council [ERC; proposal 743165] the John Innes Foundation; the JSPS [grants 15H05779, 20H05681]; and the JSPS/The Royal Society Bilateral Research for the project “Retooling rice immunity for resistance against rice blast disease” (2018-2019).

---

**Supplementary Figure 1.**
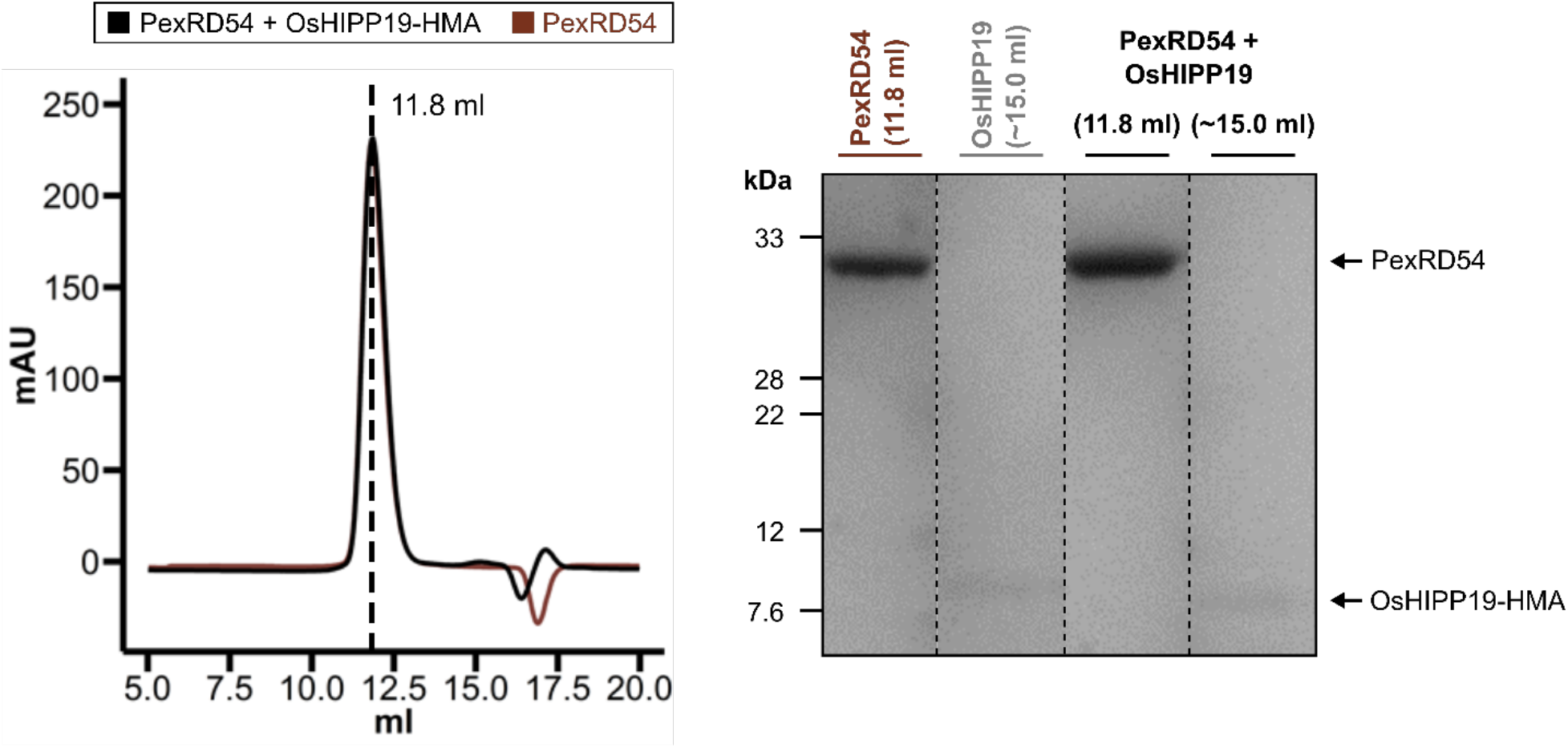
PexRD54 does not interact with OsHIPP19-HMA. Analytical gel filtration traces for PexRD54 alone (brown) and PexRD54 with OsHIPP19-HMA (black). The SDS-PAGE gel shows fractions from the peak elution volumes of each sample. No peak shift, or co-elution, is observed for these proteins.

**Supplementary Figure 2.**
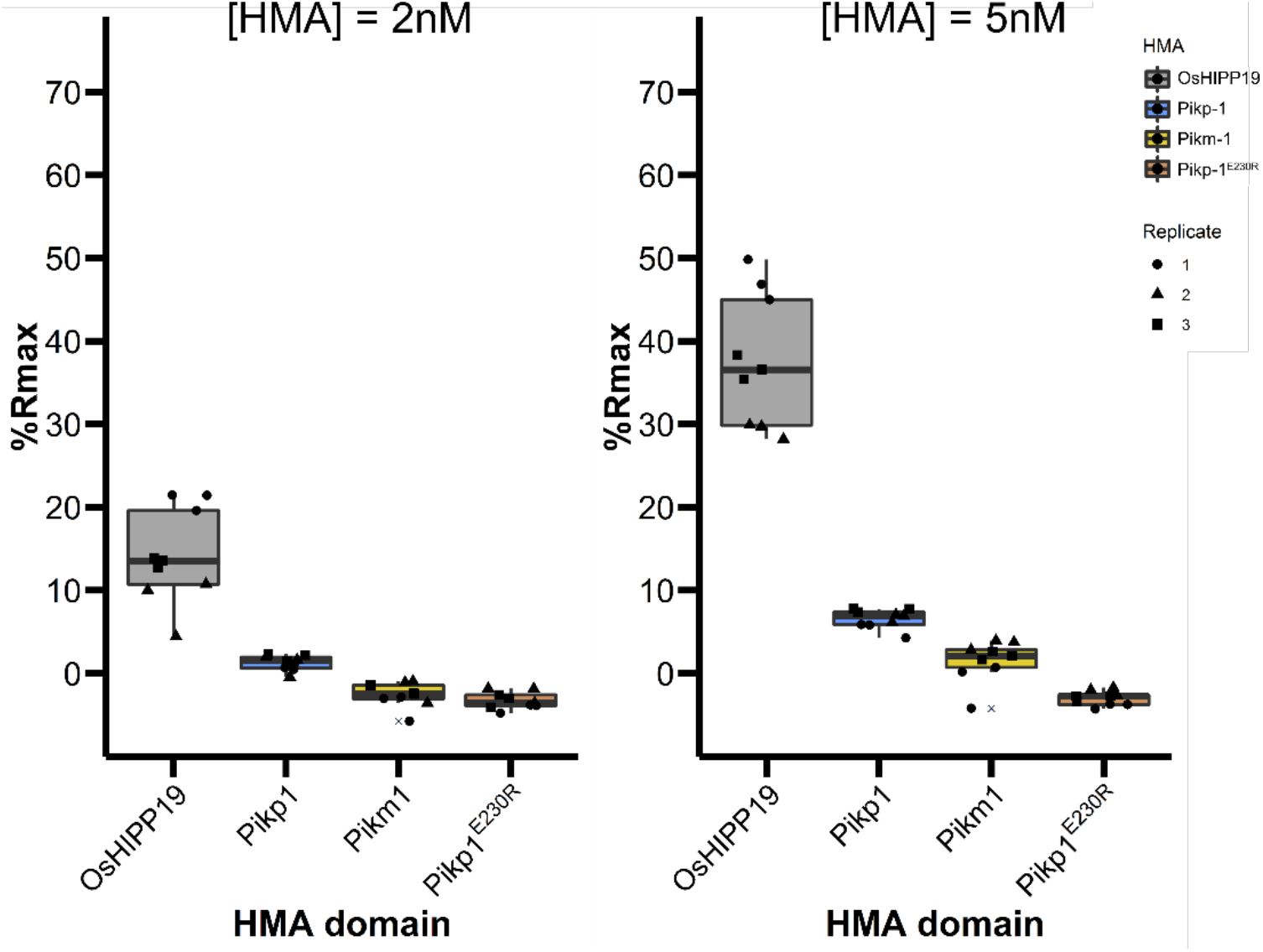
AVR-PikD interacts with the HMA domain of OsHIPP19 with higher affinity than with the integrated HMA domains of Pikp-1 or Pikm-1. %R_max_ data from two additional HMA concentrations; 2 nM (left) and 5 nM (right).

**Supplementary Figure 3.**
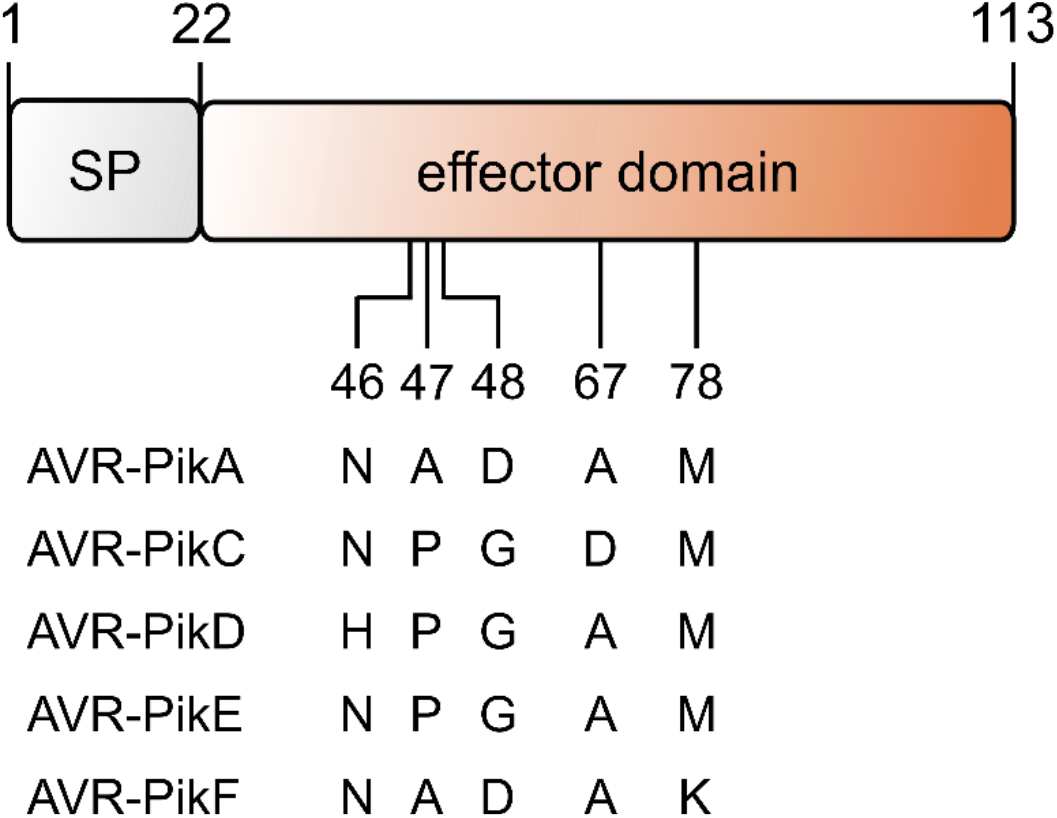
Schematic representation of the AVR-Pik effector. The identities of polymorphic amino acids for the different AVR-Pik variants are shown underneath.

**Supplementary Figure 4.**
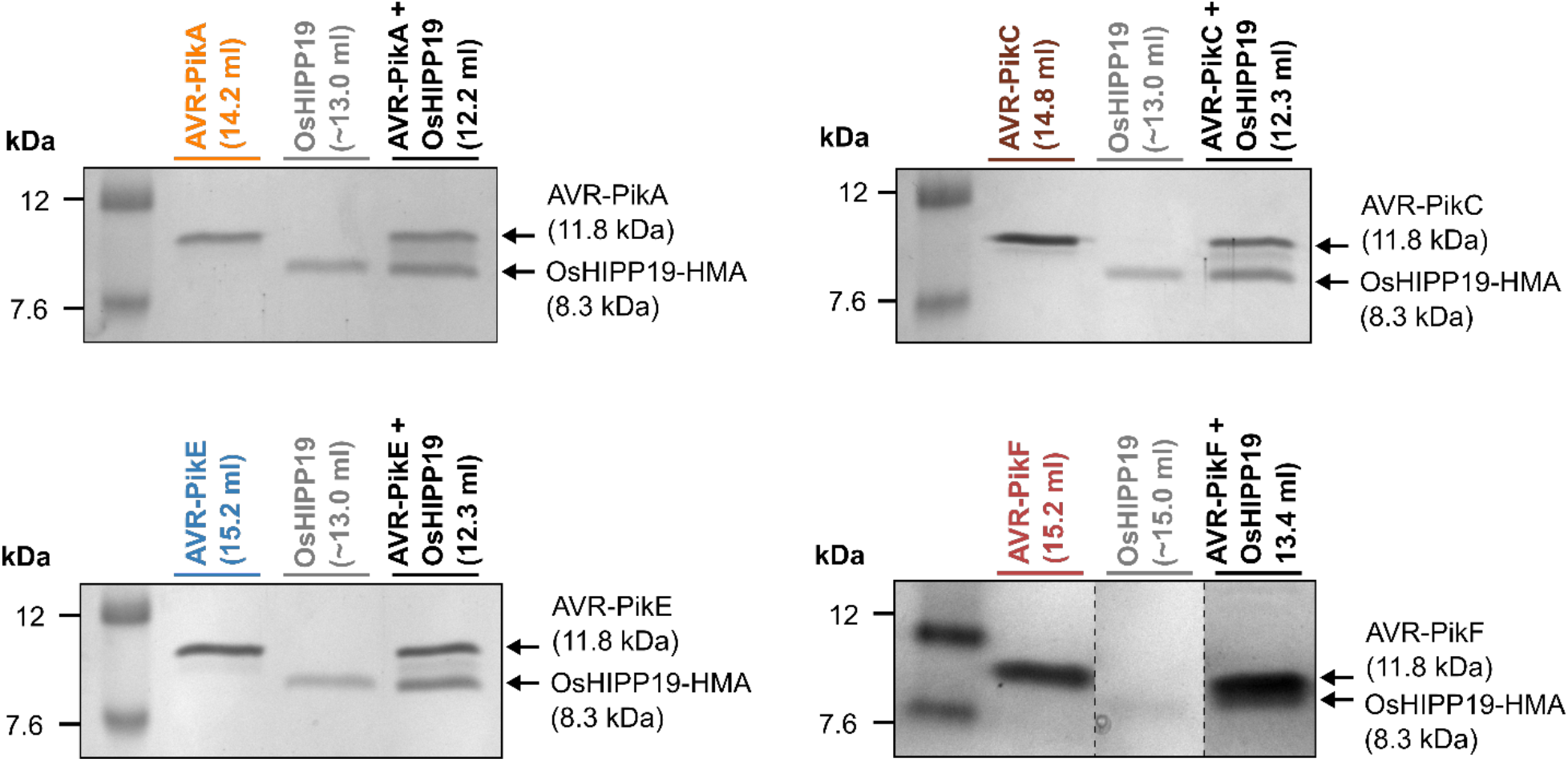
SDS-PAGE gels from analytical gel filtration experiments shown in figure 4. Gels show fractions from the peak elution volumes of each sample. The same OsHIPP19-HMA fraction was used for the gels for AVR-PikA, AVR-PikC and AVR-PikE.

**Supplementary Figure 5.**
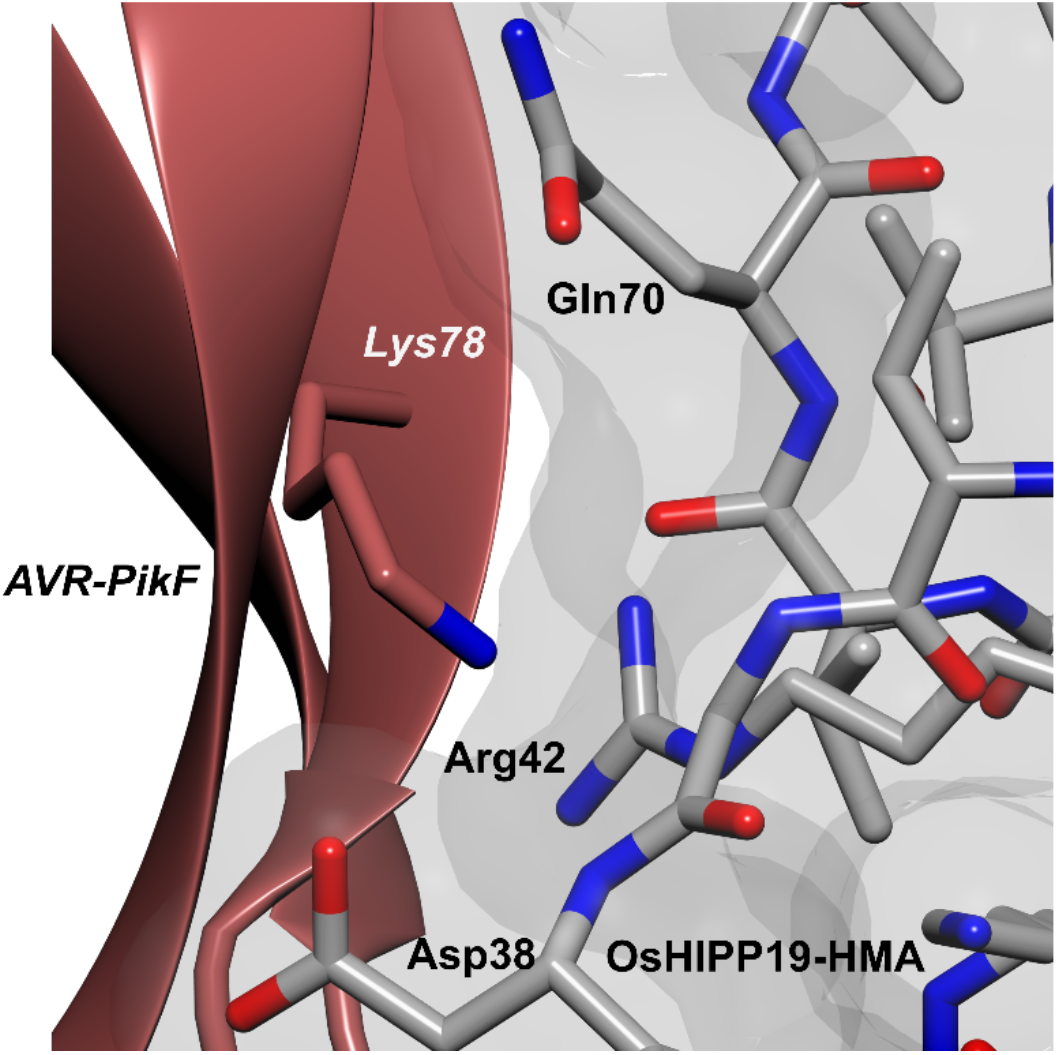
AVR-PikF^Lys78^ is located at the binding interface. The structure of AVR-PikF is represented as a red ribbon with the side chain of AVR-PikF^Lys78^ displayed as a cylinder. The structure of OsHIPP19-HMA is represented as grey cylinders with the molecular surface also displayed.

**Supplementary Table 1.**
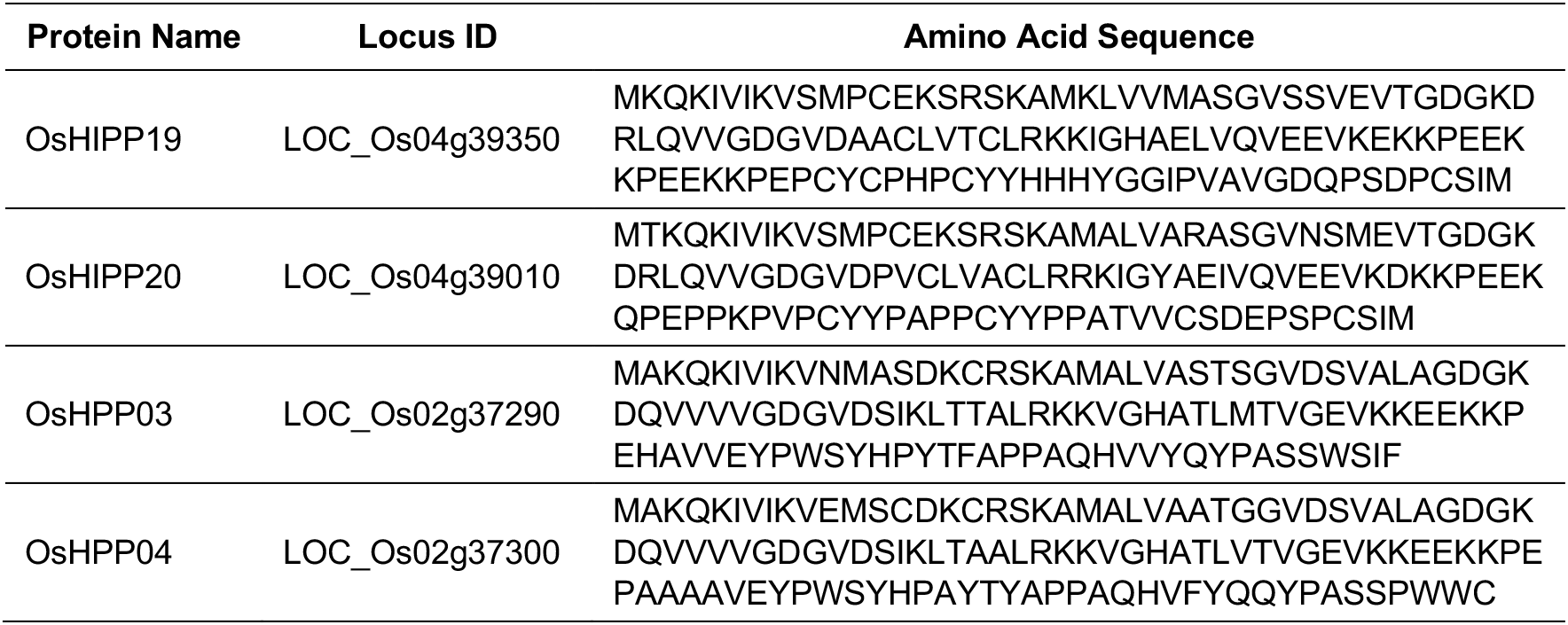
Locus IDs and amino acid sequences for the heavy metal-associated domain containing proteins referenced in this paper

**Supplementary Table 2.**
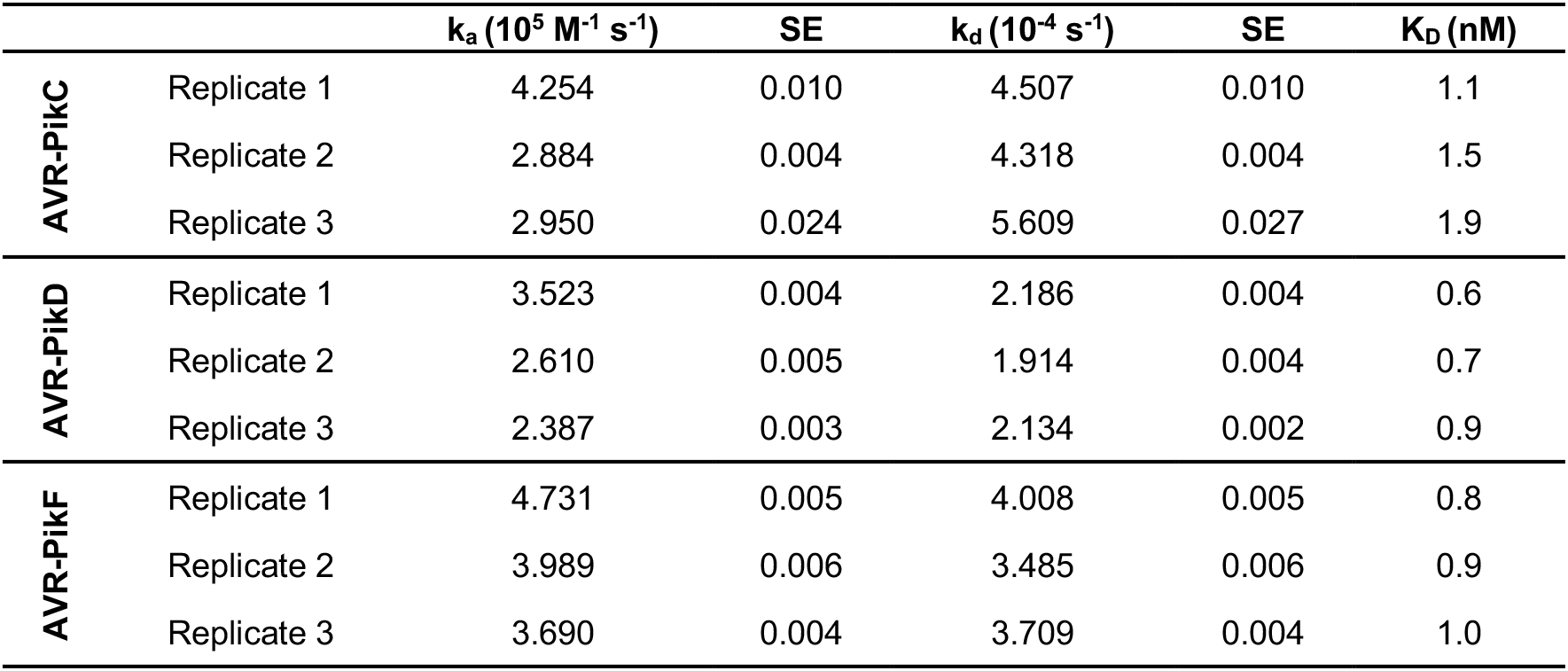
Rate constants and associated standard errors for the interactions between AVR-PikD, AVR-PikC and AVR-PikF with OsHIPP19-HMA.

| | | $k_a$ ($10^5 \text{ M}^{-1} \text{ s}^{-1}$) | SE | $k_d$ ($10^{-4} \text{ s}^{-1}$) | SE | $K_D$ (nM) |
| --- | --- | --- | --- | --- | --- | --- |
| AVR-PikC | Replicate 1 | 4.254 | 0.010 | 4.507 | 0.010 | 1.1 |
|  | Replicate 2 | 2.884 | 0.004 | 4.318 | 0.004 | 1.5 |
|  | Replicate 3 | 2.950 | 0.024 | 5.609 | 0.027 | 1.9 |
| AVR-PikD | Replicate 1 | 3.523 | 0.004 | 2.186 | 0.004 | 0.6 |
|  | Replicate 2 | 2.610 | 0.005 | 1.914 | 0.004 | 0.7 |
|  | Replicate 3 | 2.387 | 0.003 | 2.134 | 0.002 | 0.9 |
| AVR-PikF | Replicate 1 | 4.731 | 0.005 | 4.008 | 0.005 | 0.8 |
|  | Replicate 2 | 3.989 | 0.006 | 3.485 | 0.006 | 0.9 |
|  | Replicate 3 | 3.690 | 0.004 | 3.709 | 0.004 | 1.0 |

**Supplementary Table 3.** Comparison of the binding interfaces of OsHIPP19-HMA/AVR-PikF and Pikm-HMA/AVR-PikA complexes. Interface analysis was performed using qtPISA (70).

| Interface parameter | OsHIPP19-HMA /<br>AVR-PikF | Pikm-HMA /<br>AVR-PikA |
| --- | --- | --- |
| Interface area (Å<sup>2</sup>) | 1045.3 | 918.2 |
| Solvation energy (kcal/mol) | 0.1 | -5.6 |
| Binding energy (kcal/mol) | -11.2 | -13.5 |
| Hydrophobic P-value | 0.6917 | 0.4405 |
| Hydrogen bonds | 17 | 12 |
| Salt bridges | 10 | 7 |
| Disulphide bonds | 0 | 0 |

